# Dynamic PET of Human Liver Inflammation: Impact of Kinetic Modeling with Optimization-Derived Dual-Blood Input Function

**DOI:** 10.1101/268748

**Authors:** Guobao Wang, Michael T. Corwin, Kristin A. Olson, Ramsey D. Badawi, Souvik Sarkar

## Abstract

The hallmark of nonalcoholic steatohepatitis is hepatocellular inflammation and injury in the setting of hepatic steatosis. Recent work has indicated that dynamic ^18^F-FDG PET with kinetic modeling has the potential to assess hepatic inflammation noninvasively, while static FDG-PET did not show a promise. Because the liver has dual blood supplies, kinetic modeling of dynamic liver PET data is challenging in human studies. This paper aims to identify the optimal dual-input kinetic modeling approach for dynamic FDG-PET of human liver inflammation. Fourteen patients with nonalcoholic fatty liver disease were included. Each patient underwent 1-hour dynamic FDG-PET/CT scan and had liver biopsy within six weeks. Three models were tested for kinetic analysis: traditional two-tissue compartmental model with an image-derived single-blood input function (SBIF), model with population-based dual-blood input function (DBIF), and new model with optimization-derived DBIF through a joint estimation framework. The three models were compared using Akaike information criterion (AIC), F test and histopathologic inflammation score. Results showed that the optimization-derived DBIF model improved liver time activity curve fitting and achieved lower AIC values and higher F values than the SBIF and population-based DBIF models in all patients. The optimization-derived model significantly increased FDG K_1_ estimates by 101% and 27% as compared with traditional SBIF and population-based DBIF. K_1_ by the optimization-derived model was significantly associated with histopathologic grades of liver inflammation while the other two models did not provide a statistical significance. In conclusion, modeling of DBIF is critical for dynamic liver FDG-PET kinetic analysis in human studies. The optimization-derived DBIF model is more appropriate than SBIF and population-based DBIF for dynamic FDG-PET of liver inflammation.

## INTRODUCTION

Nonalcoholic fatty liver disease (NAFLD) affects approximately 30% of the general population and is emerging as a leading cause of liver-related morbidity and mortality (*1,2*). Nonalcoholic steatohepatitis (NASH) is a more severe form of NAFLD characterized by hepatocyte inflammation and injury and can subsequently lead to cirrhosis and associated liver cancer and liver failure (*3*). NASH develops in 5-10% of NAFLD patients (i.e., 5-10 million people in the United States) and is associated with higher liver-related mortality than is hepatic steatosis alone (*4*). Differentiation of NASH from simple fatty liver is essential for future patient management in NAFLD, particularly as newer therapies become available (*3,5*).

The diagnostic hallmark of NASH is hepatocellular inflammation and injury in the setting of hepatic steatosis (*3*). Imaging methods have been developed to quantify hepatic steatosis (*6*), e.g. ultrasonography (*7*), computed tomography (CT) (*8*) or magnetic resonance imaging (MRI) (*9–11*). Other methods such as magnetic resonance elastography (MRE) (*12*) and ultrasound elastography (*13*) can measure liver stiffness and quantify advanced fibrosis of hepatic tissue (*14–16*). However, there is currently no effective imaging method for accurate characterization of liver inflammation in clinical practice and clinical trials (*17*). Liver biopsy followed by clinical histopathology remains the current standard practice (*18*).

^18^F-fluorodeoxyglucose (FDG) positron emission tomography (PET) is a widely used and effective method for imaging cell glucose metabolism. Previous studies have reported the use of FDG-PET for imaging infection and inflammation (*19,20*). Several studies have investigated the use of FDG-PET in the liver (*21–27*). However, none of these studies have reported relevant or promising results for liver inflammation and NASH assessment. It is worth noting that current clinical usage of PET is largely limited to static PET imaging which provides standardized uptake value (SUV) as a semi-quantitative measure of glucose utilization. This static way of using PET may not be able to explore the actual role of FDG-PET in NAFLD and NASH. Our recent preliminary work has indicated that while SUV by static FDG-PET did not show a promise, dynamic FDG-PET with kinetic modeling is promising for assessing liver inflammation in NASH (28). The FDG K_1_ parameter, which represents the transport rate of FDG from blood to hepatic tissue, can be a potential PET biomarker for characterizing liver inflammation.

Blood input function is essential for kinetic modeling in dynamic FDG-PET (*29*). Conventionally it is either obtained by invasive arterial blood sampling or noninvasively derived from the left ventricle or aortic regions in dynamic PET images (*30–32*). In the liver, both hepatic artery and portal vein provide blood supply to the hepatic tissue (*22*). The portal vein input is not the same as the hepatic artery input and is difficult to obtain in human objects (*22*). Neglecting the influence of portal vein can result in inaccurate kinetic modeling. A common approach to accounting for the effect of dual blood supplies is to use a flow-weighted model of dual-blood input function (DBIF) (*21,33,34*). The model parameters are often pre-determined by population means that were derived using arterial blood sampling in animal studies (*22,35*). Due to heterogeneity across patients and potential differences between arterial and image-derived input functions, the population-based DBIF model may become ineffective when arterial blood sampling is not available and image-derived input function is actually used in human studies.

The objective of this paper is to evaluate and identify an appropriate dual-input kinetic modeling approach for FDG kinetic quantification of liver inflammation using human patient data. Besides the population-based DBIF approach, we also examined an optimization-derived DBIF model which employs mathematical optimization to jointly estimate the parameters of DBIF and liver FDG kinetics. This model directly utilizes image-derived aortic input function, requires no arterial blood sampling, and is more adaptive to individual patients. We used both statistical information criteria and histopathologic inflammation data to compare these methods.

## MATERIALS AND METHODS

### Patient Characteristics

Fourteen patients with NAFLD were included in this study. These patients had a liver biopsy as a part of routine clinical care or for enrollment in clinical trials. The University of California Davis Institutional Review Board and the University of California Davis Medical Center Radiation Use Committee approved the study. All patients signed informed consent before participating in the study. Patients with history of alcohol abuse, chronic hepatitis B or C, or other chronic liver disease other than NAFLD were excluded from the study.

### Liver Histopathology

Liver biopsies were performed under ultrasound guidance and scored according to the nonalcoholic steatohepatitis clinical research network (NASH-CRN) criteria (*36*). The NAFLD activity score (NAS, range 0-8) is comprised of severity of steatosis, inflammation, and hepatocellular ballooning. While a NAS score greater than 4 has been reported to correlate with the presence of NASH, the scores of lobular inflammation and ballooning degeneration are noted to represent hepatic inflammation and injury, and are therefore combined to create an overall “hepatic inflammation” score (range 0-5). In this study, an inflammation score >3 was considered indicative of high inflammation, and a score <3 and score =3 were deemed as low and medium inflammation, respectively.

### Dynamic ^18^F-FDG PET/CT

#### Scan Protocol

Dynamic ^18^F-FDG PET studies were performed with the GE Discovery 690 PET/CT scanner at the UC Davis Medical Center. Patients fasted for at least 6 hours. Each patient was injected with 10 mCi ^18^F-FDG. List-mode time-of-flight data acquisition started right after the intravenous bolus administration and lasted for one hour. At the end of PET scan, a low-dose CT scan was performed for attenuation correction for PET. Dynamic PET data were binned into 49 time frames using the sampling schedule: 30 ⨯ 10s, 10 ⨯ 60s, and 9 ⨯ 300s. Dynamic PET images were then reconstructed using the vendor software with the standard ordered subsets expectation maximization (OSEM) algorithm with 2 iterations and 32 subsets.

#### Extraction of Blood and Tissue Time Activity Curves (TACs)

Eight spherical regions of interest (ROI), each with 25 mm in diameter, were placed on the eight segments of the liver avoiding any major blood vessels. These ROI placements were tuned and confirmed by an abdominal radiologist. The averaged FDG activity in all 8 ROIs was extracted from the dynamic images to form a global liver TAC of low noise. An additional ROI is placed in the descending aorta region to extract image-derived aortic input function.

**FIGURE 1:**
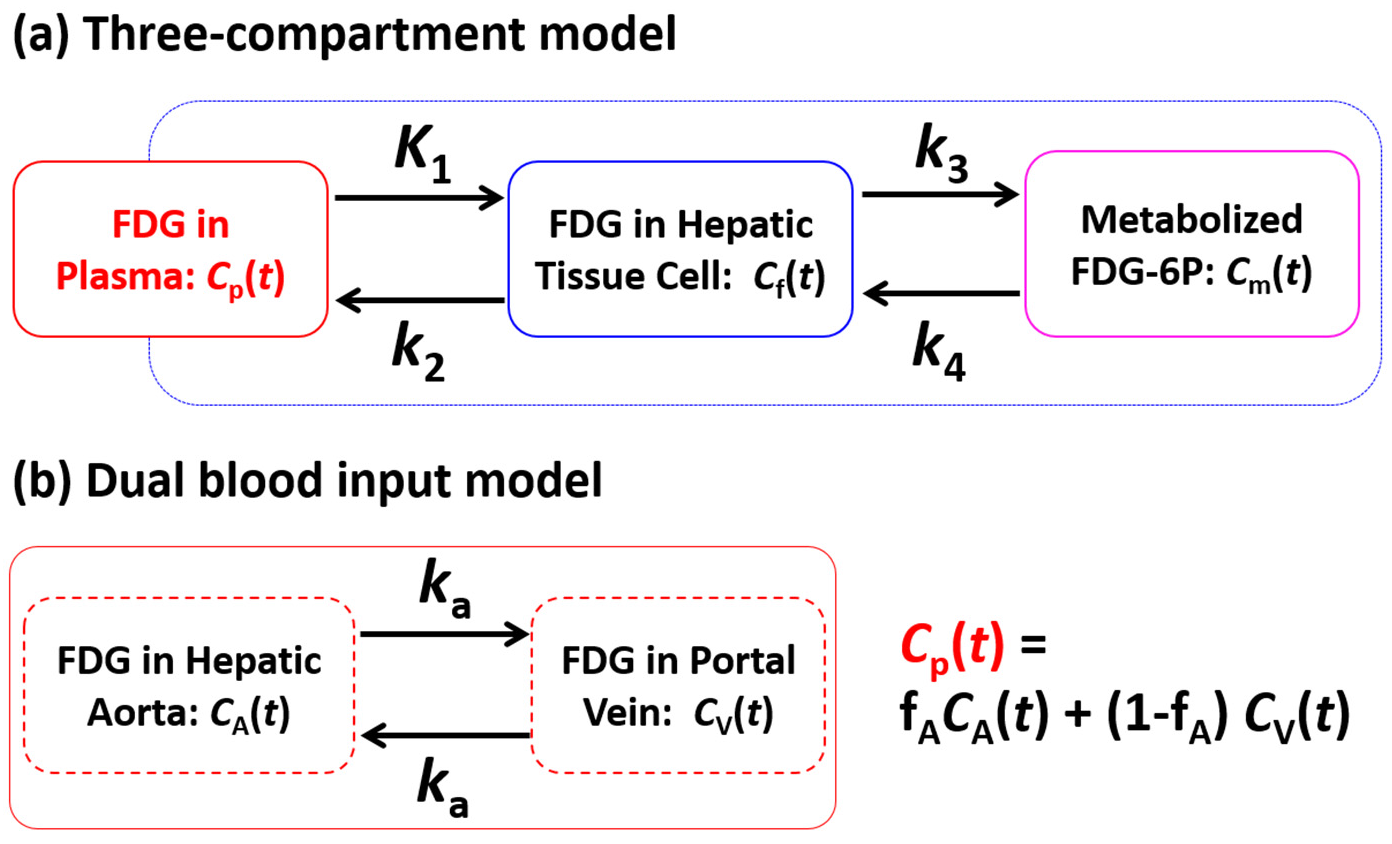
Kinetic modeling of dynamic liver FDG-PET data with a dual-blood input function from the hepatic artery and portal vein.

### Dual-Input Kinetic Modeling

#### Compartmental Model with Dual-blood Input Function (DBIF)

FDG kinetics commonly follow the two-tissue compartmental model (*29*) as shown in Figure 1(a). Glucose transporters transport ^18^F-FDG from blood to hepatic tissue with the rate constant *K*_1_ (mL/min/mL) and from hepatic tissue to blood with the rate *k*_2_ (1/min). FDG is phosphorylated by hexokinase in cells into FDG 6-phosphate with the rate *k*_3_ (1/min) and the dephosphorylation process occurs with the rate *k*_4_ (1/min). *C*_*p*_(*t*), *C*_*f*_(*t*) and *C*_*m*_(*t* represent the FDG concentration in the plasma compartment, free-state FDG in the hepatic tissue compartment, and metabolized FDG 6-phosphate in the tissue, respectively. In traditional kinetic modeling with single-blood input function (SBIF), only blood supply from the aorta is considered and thus the input function *C*_*p*_(*t*) is equivalent to the aortic input function *C*_*A*_(*t*). To account for the effect of dual blood supplies in the liver, a flow-weighted sum of the aortic input *C*_*A*_(*t*) and portal vein input *C*_*v*_(*t*) can be used to model the DBIF (*22*): 
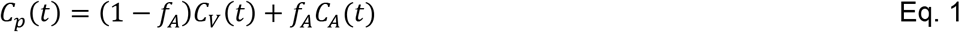
where *f*_*A*_ is the fraction of hepatic artery contributing to the overall liver blood flow. As shown in Figure 1(b), portal vein can be considered as an additional compartment given FDG in the portal vein flows through the gastrointestinal system with the rate *k*_*a*_ (1/min) before entering into the liver. Thus, the portal vein input function *C*_*v*_(*t*) follows an analytical solution 
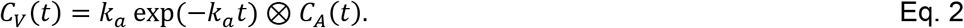

This is equivalent to a dispersed version of the aortic input function *C*_*A*_(*t*) (*37*). As a result, the combined compartmental model is equivalently described by a set of differential equations: 
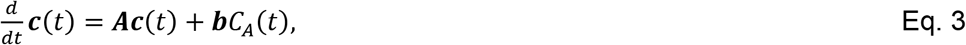
where 
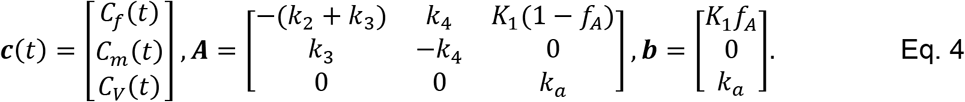

The total activity that can be measured by PET is the sum of different compartments:

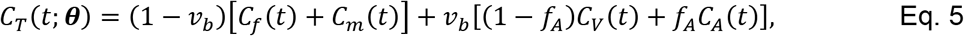

where *υ*_*b*_ is the fractional blood volume and *θ* = [*υ*_*b*_,*K*_1_,*k*_*2*_,*k*_3_,*k*_4_,*k*_*a*_,*f*_*A*_]^*T*^. is a vector collecting all unknown parameters. Note that this DBIF model becomes the traditional SBIF model when *k*_*a*_ = 0 and *f*_*A*_ = 1.

#### Joint Estimation of Kinetic and Input Parameters

All model parameters are jointly estimated by fitting a measured liver TAC č using the model equation and following nonlinear least square estimation:

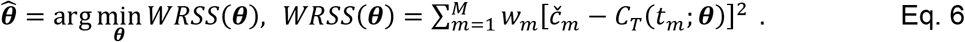

where *WRSS*(*θ*) denotes the weighted residual sum of squares of the curve fitting and *w*_*m*_ denotes the weighting factor for time frame *m.* We used the classic Levenberg-Marquardt algorithm to solve the optimization problem. In this paper, we refer this joint estimation as the optimization-derived DBIF approach for clarity. It would become equivalent to the population-based DBIF if the parameters *f*_*A*_ and *k*_*a*_ are assigned with fixed population means (if known). Based on our initial analysis of the patient data sets, we set the initials of kinetic parameters to *υ*_*b*_ = 0.01, *k*_1_ = 1.0, *k*_*2*_ = 1.0, *k*_3_ = 0.01, *k*_4_ = 0.01, *k*_*a*_ = 1.0, *f*_*A*_ = 0.01.

### Comparison of Kinetic Models

We compared three input models: traditional model with SBIF, model with population-based DBIF and model with optimization-derived DBIF. In this study, the SBIF was derived from the descending aorta region in dynamic FDG-PET images. To utilize reported population means for the population-based DBIF, we used the following portal vein input model (*22*)

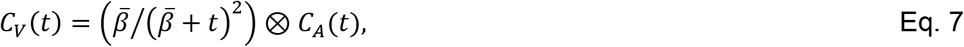

which is very similar to the exponential-based model in Eq. 2 but not easily integrated into the differential equations for joint estimation. 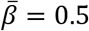 0.5 and *f*_*A*_ = 0.25 were previously reported for FDG (*22*).

#### Comparison Using Statistical Criteria

The three models were compared using two established statistical criteria for model selection for TACs: corrected Akaike information criterion (AIC) and F test (*38,39*). The AIC is defined by

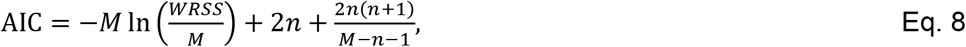

where *n* denotes the total number of unknown parameters. Note that *n*=5 for the SBIF and population-based DBIF models and *n*=7 for the optimization-derived DBIF model. Here AIC was corrected for finite sample sizes due to the ratio 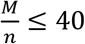. A lower AIC value indicates a better selection of model.

The F test compares a complex model 2 with a simple model 1 using

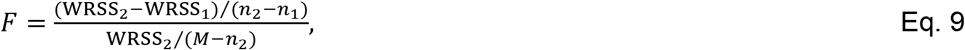

where the degree of freedom (number of unknown parameters) *n*_2_<*n*_1_. A larger F value indicates better fit. If the p value of F test is 0.05 or less, the model 2 is then considered significantly better than model 1.

#### Evaluation Using Histopathological Inflammation Data

Patients in this study were divided into three groups according to their histopathological inflammation scores: low inflammation (<3), medium inflammation (=3), and high inflammation (>3). To examine the capability of FDG kinetic parameters for differentiating different inflammation groups, a two-sample 2-sided t-test at the 0.05 level was used to test the difference of group means and the Mann-Whitney U test was used to test difference of group medians. All statistical analyses were conducted using MATLAB (Natick, MA). P-values less than 0.05 were considered as statistically significant in this study.

## RESULTS

### Patient Characteristics

Majority of the patients were white (75%) while 25% were Hispanics. Female patients formed 67% of the cohort with 75% of the patients between the ages 40-70 years and 25% between 18-39 years. The mean BMI was 34±6 kg/m^2^. The mean fasting glucose prior to PET scan was 115±33 mg/dL. The patient population had an equitable spread across NAS score (≥5 of 58%). Two thirds of patients had hepatic inflammation sore of ≥3.

### Demonstration of dual-blood input function

Figure 2 demonstrates an optimization-derived DBIF from a patient data set. Only the first 5 minutes are shown in the figure and the curves after 5 minutes are very similar among others. The DBIF is a weighted combination of the image-derived SBIF and the portal vein input function derived from the optimization. In this example, the weighting factor *f*_*A*_ was 0.19. The DBIF has a much lower peak value than the SBIF because the contribution from hepatic artery is small (19%) while the contribution from portal vein is great (81%).

**FIGURE 2:**
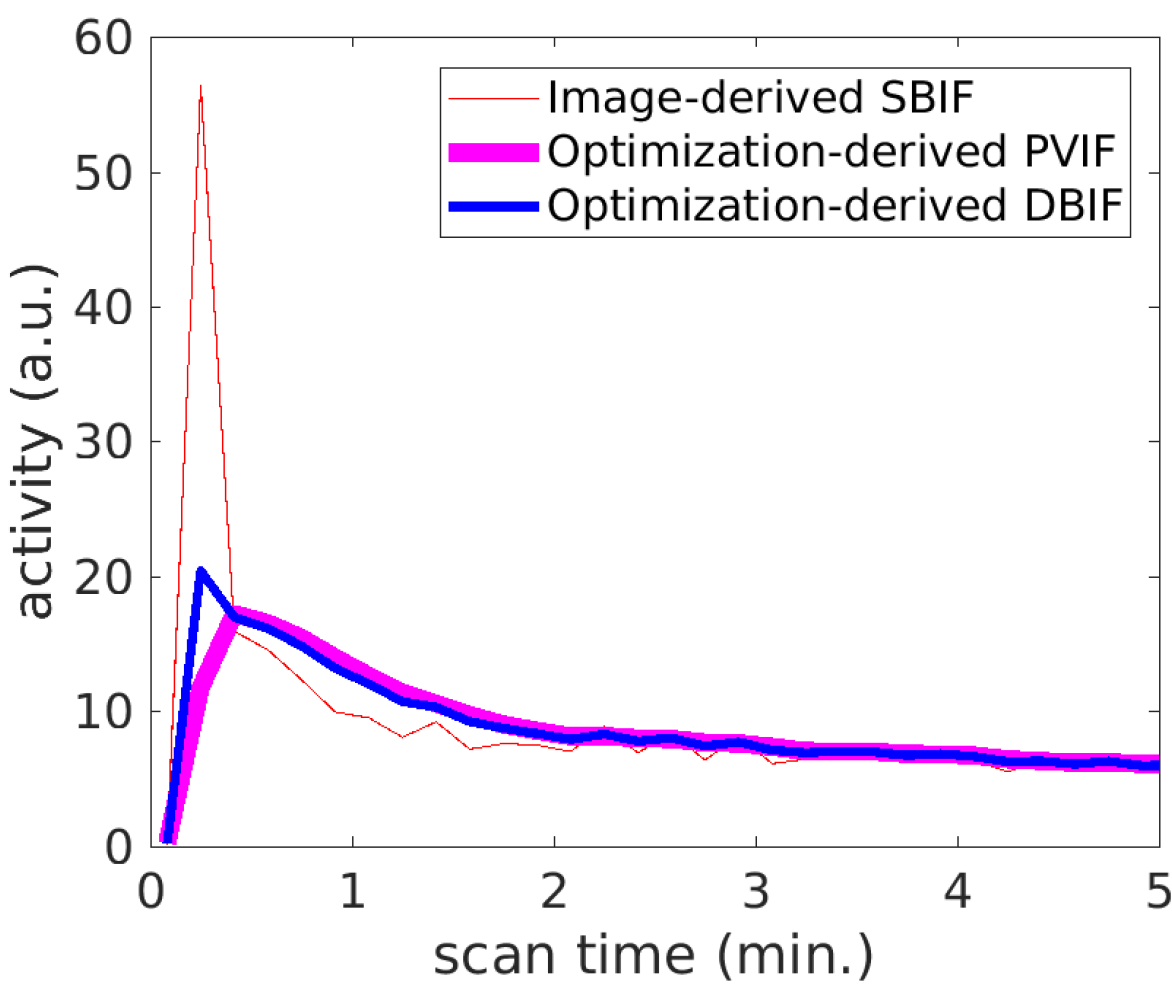
Example of optimization-derived DBIF and its components from hepatic artery and portal vein: image-derived SBIF and portal vein input function (PVIF).

**FIGURE 3:**
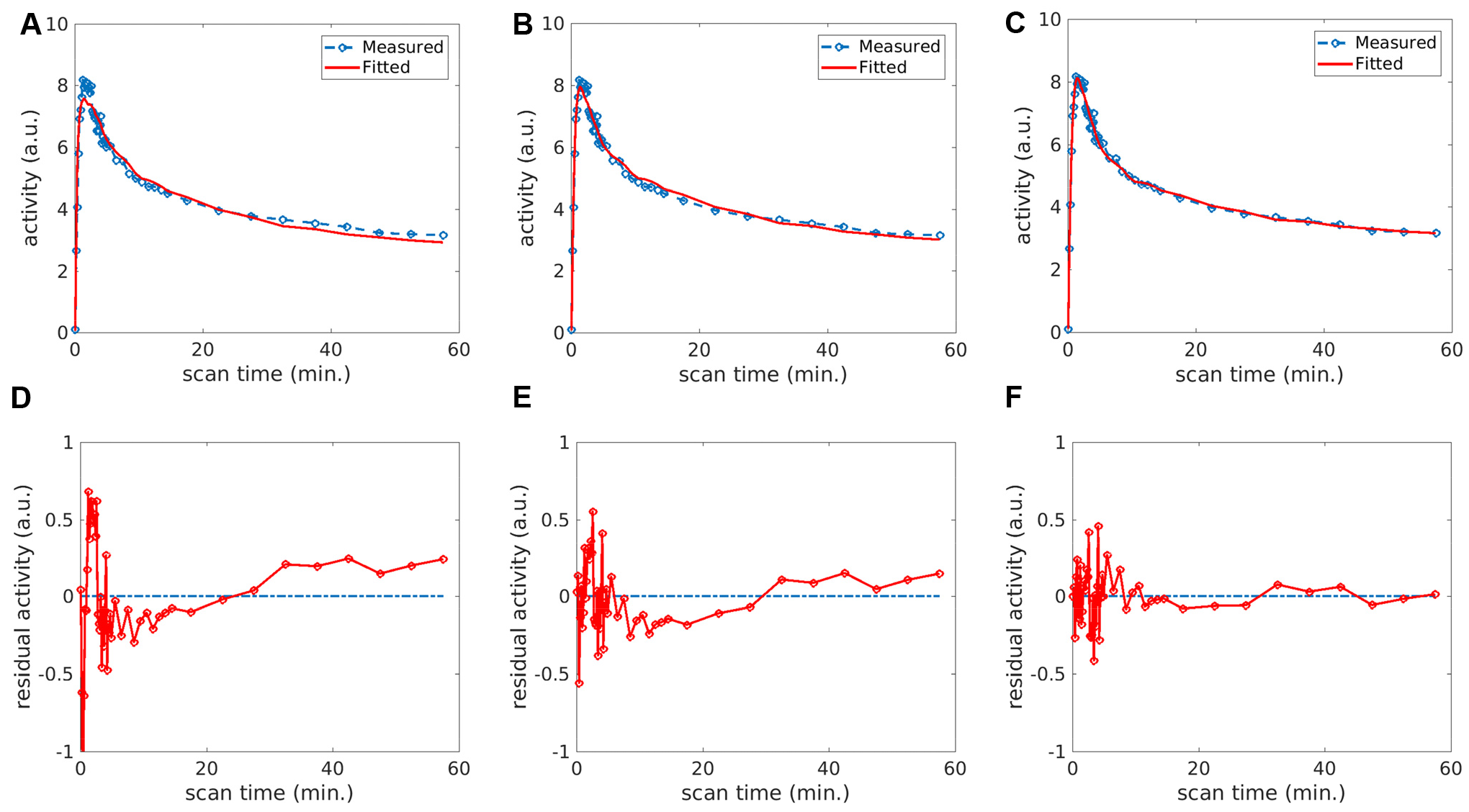
Fit of a liver TAC using different input models: (A) SBIF, (B) population-based DBIF, (C) optimization-derived DBIF; (D-F) corresponding residual plots of the fits in (A-C).

**FIGURE 4:**
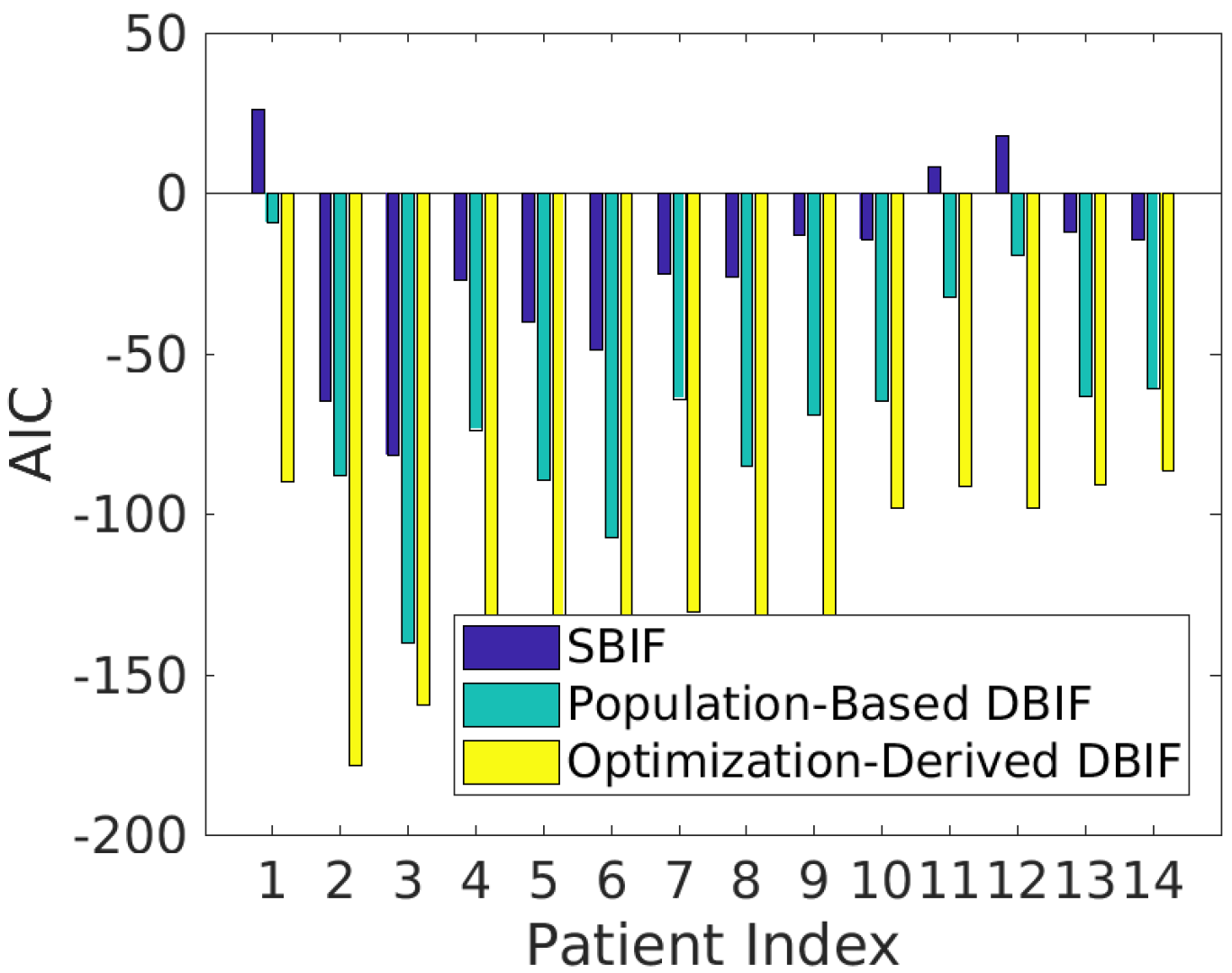
AIC values of liver TAC fitting with different input function models.

### Evaluation of TAC Fit Quality

Figure 3A-3C show the fittings of the liver TAC using the image-derived SBIF, population-based DBIF and optimization-derived DBIF. The residual plots of these fits are shown in Figure 3D-3F. The fitting with SBIF could not fit the early-time peak and late time points due to lack of the portal vein component in the input function. The population-based DBIF provided an improved fit of the peak but still suffered error for late time points due to inaccuracy of the population means for a specific patient data set. A linear trend was observed in the residual plots (Figure 3D and 3E) for these two approaches, indicating a systematic bias in the fitting. The optimization-derived DBIF estimated the input parameters from the data and fit both the peak at early time and late time points closely. The linear trend was removed and the residuals became asymptotically normal.

The AIC values of the three fittings were −81, −140 and −159, respectively. AIC values of the three input models are further compared in Figure 4 for all individual patients in the cohort. The average AIC was −23±30 by SBIF, −69±34 by the population-based DBIF, and −128±34 by the optimization-derived DBIF. The optimization-derived DBIF model had the lowest AIC in all patients.

The results of F test are given in Table 1. The minimum F value among different patients was 80.5 for comparing the optimization-derived DBIF with SBIF and 13.4 for comparing the optimization-derived DBIF with population-based DBIF model, both greater than the F critical value 3.2 calculated with *n*_1_ = 5 and *n*_2_ = 7 for a p value of 0.05. The p values of F test in individual patients are all small (<0.0001), indicating the optimization-derived DBIF is more appropriate for TAC fitting than the traditional SBIF and population-based DBIF models.

### Change in Kinetic Parameters

The three input models resulted in different estimates in kinetic parameters. The mean and standard deviation of kinetic parameters *K*_1_,*k*_2_,*k*_3_,*k*_4_ and *K*_*i*_ = *K*_1_ *k*_3_/(*k*_2_ + *k*_3_) estimated by the three approaches are listed in Table 2. It is worth noting that a reversible kinetic model (i.e., *k*_4_ > 0) was required in our study for accurately modeling FDG TACs in the liver. Neglecting *k*_4_ in the model reduced TAC fit quality with higher AIC values (results not shown). Compared with the image-derived SBIF, the optimization-derived DBIF significantly increased the mean values of *K*_1_(0.5112 vs 0.9829), *k*_2_ (0.4983 vs 1.1053), *k*_3_ (0.0008 vs 0.0141) and *K*_*i*_ (0.0008 vs 0.0119), all with p<0.0001. The difference in *K*_1_ in individual patient was 101% on average and up to 150%, as shown in Figure 5. Compared with the population-based DBIF, the optimization-derived DBIF significantly increased the mean values of *k*_3_ (0.0011 vs 0.0141, p<0.0001), *k*_4_ (0.0196 vs 0.0534, p=0.0034) and *K*_*i*_ (0.0010 vs 0.0119, p<0.0001). Although the mean values of by the optimization-derived DBIF and population-based DBIF models are similar to each other (0.9787 vs. 0.9829), *K*_1_ values by the two models were very different in each individual patient (Figure 5). The change was 27% on average and up to 44%.

**TABLE 1:** F statistics of model comparison (F critical value is 3.2 at the significance level of 0.05)

| Model Comparison | F Values |  |  | P Values |
| --- | --- | --- | --- | --- |
| | Mean±SD | Min | max | |
| Optimization-Derived DBIF vs SBIF | 200.2±102.7 | 80.5 | 389.9 | <0.0001 |
| Optimization-Derived DBIF vs Population-Based DBIF | 67.5±43.7 | 13.4 | 137.7 | <0.0001 |

**TABLE 2:** Mean and standard deviation of FDG kinetic parameters estimated by different input function approaches

| | $K_1$ | $k_2$ | $k_3$ | $k_4$ | $K_i$ |
| --- | --- | --- | --- | --- | --- |
| SBIF | 0.5112±0.2064 | 0.4983±0.1760 | 0.0008±0.0010 | 0.0512±0.0465 | 0.0008±0.0010 |
| Population-Based DBIF | 0.9787±0.5509 | 0.9197±0.4222 | 0.0011±0.0011 | 0.0196±0.0291 | 0.0010±0.0011 |
| Optimization-Derived DBIF | 0.9829±0.2730 | 1.1053±0.2488 | 0.0141±0.0070 | 0.0534±0.0263 | 0.0119±0.0051 |

**FIGURE 5:**
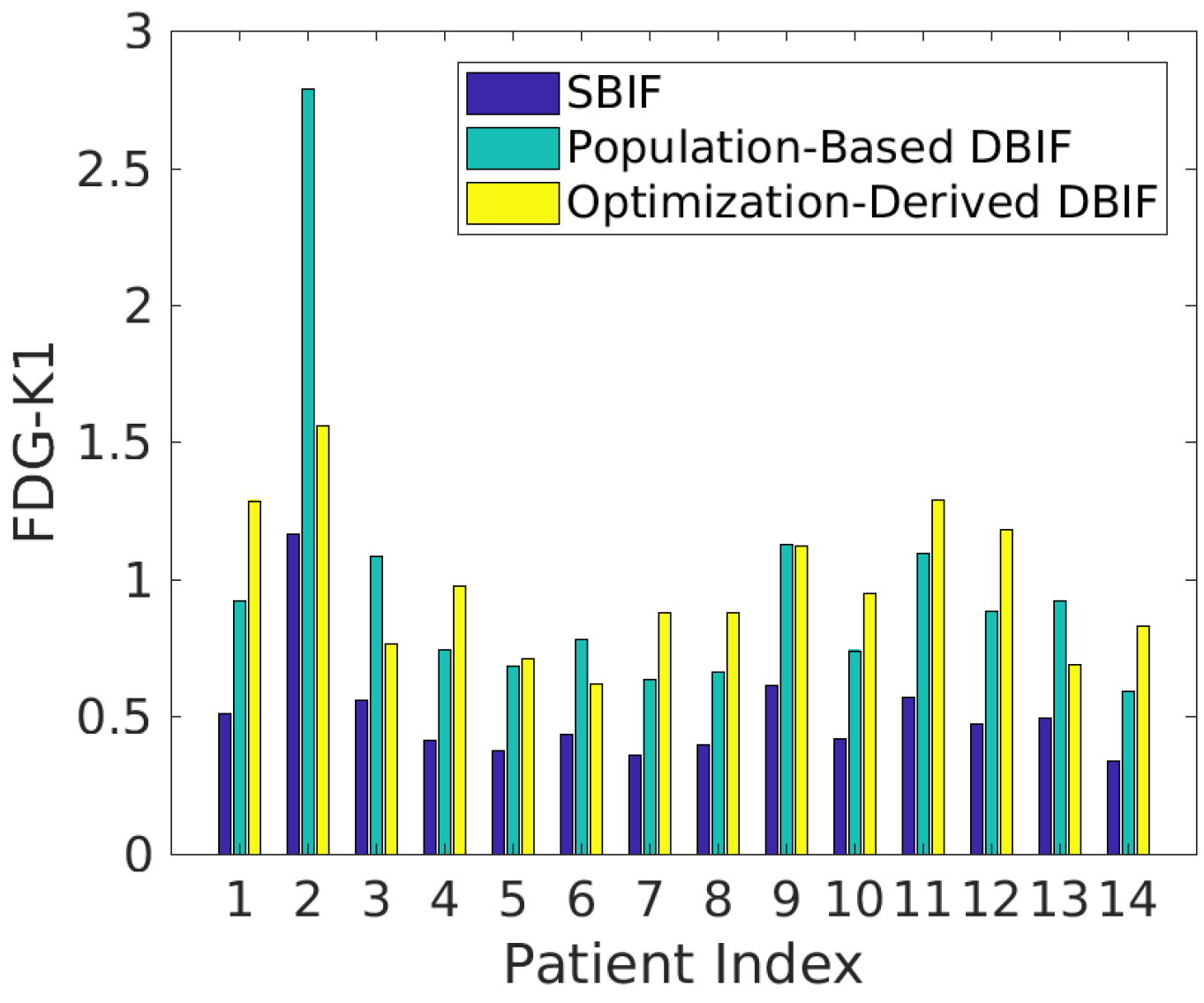
FDG Ki values estimated by different input function models.

**FIGURE 6:**
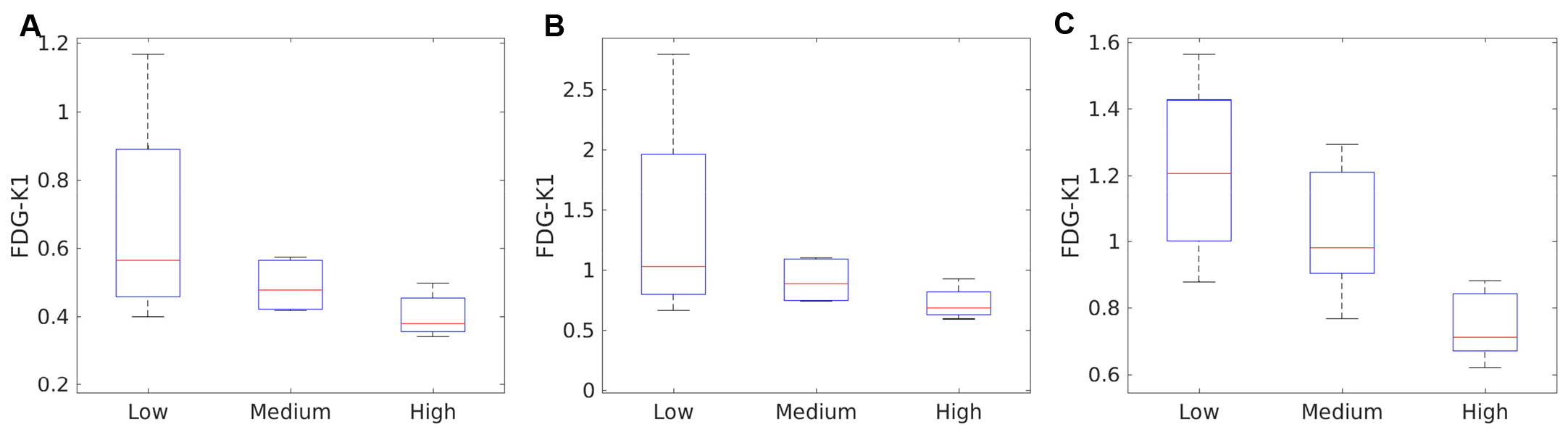
Association of histopathological inflammation score with FDG-K_1_ estimated by different input models. (A) Image-derived SBIF; (B) Population-based DBIF; (C) Optimization-derived DBIF.

**TABLE 3:** P values of t test and U test for comparing in different liver inflammation groups using FDG K_1_ estimated by SBIF, population-based (PB) DBIF, and optimization-derived (OD) DBIF.

| Group Comparison | T test |  |  | U test |  |  |
| --- | --- | --- | --- | --- | --- | --- |
|  | SBIF | DBIF-PB | DBIF-OD | SBIF | DBIF-PB | DBIF-OD |
| Low vs. medium inflammation | 0.2733 | 0.3158 | 0.3121 | 0.5556 | 0.5556 | 0.5556 |
| Medium vs. high inflammation | 0.0826 | 0.0941 | **0.0242** | 0.1508 | 0.1508 | **0.0317** |
| Low vs. high inflammation | 0.1206 | 0.1707 | **0.0116** | 0.0635 | 0.1905 | **0.0317** |

### Association with Histopathologic Data

Figure 6 shows the associations of histopathologic grades of liver inflammation with FDG K_1_ by different input models. The standard boxplots reflect the median, 25% percentile and 75% percentile of K_1_ in each of low, medium and high inflammation groups. Overall, FDG K_1_ value decreased as inflammation grade increased. The results of statistical t test for comparing group means and U test for comparing group medians are summarized in Table 3. Neither the SBIF model nor the population-based DBIF model could differentiate low, medium and high inflammation groups (p>0.05). In comparison, K_1_ by the optimization-derived DBIF model was better associated with the inflammation grades and differentiated low versus high inflammation groups and medium versus high inflammation groups (p<0.05).

## DISCUSSION

Determination of liver inflammation in nonalcoholic fatty liver disease patients is crucial in differentiating serious NASH from simple hepatic steatosis. Dynamic FDG-PET with kinetic modeling has the potential to provide a noninvasive imaging biomarker for characterizing hepatic inflammation. Accurate kinetic modeling of dynamic liver FDG-PET data requires consideration of the effect of dual-blood supplies in the liver. Although the input function from the hepatic artery can be derived from the aorta in the dynamic images, it is difficult to derive the portal vein input function from dynamic PET because the size of portal vein (10-15mm) is small compared to the spatial resolution of clinical PET scanner (4-8mm). Partial volume effects, noise and respiratory motion all can contaminate the accuracy of the portal vein input function.

The standard SBIF model simply neglects the portal vein input and therefore provides inaccurate estimation of FDG kinetic parameters. The K_1_ parameter, which is the major parameter of interest for liver inflammation, is generally underestimated by SBIF, as demonstrated in this study. To account for the dual-blood effect, we applied the population-based DBIF model (*21,33*) in this study. It improved TAC fitting with lower AIC and higher F values than SBIF but did not improve association of the FDG biomarker with inflammation grades. This can be explained by the fact that the population model is not adaptive to individual objects and thus might not provide the optimal estimation. In addition, the aortic input function was derived from PET images while the population-based model was derived using arterial sampling, resulting in further suboptimal performance.

The optimization-derived DBIF model provided the best performance according to TAC fitting quality and association with histopathological inflammation grades. Because the parameters of the input model were jointly estimated with liver tissue kinetic parameters, the estimation was more adaptive to individual patients and achieved lower AIC and higher F values. In this study, the two DBIF model parameters k_a_ and f_A_ were 1. 627±1.427 and 0.044±0.054. Here the f_A_ estimates were far smaller than the population mean 0.25 that was reported based on arterial sampling. This is possibly explained by the difference between image-derived input function and arterial blood input. The resulting K_1_ parameter estimates were significantly different from those by SBIF and optimization-based DBIF. Although there was no ground truth of K_1_ values for validating the estimates, evaluation of statistical information criteria and analysis of association of K_1_ with liver inflammation grades provided a feasible way to prove the improved performance of the modified DBIF model.

In addition to establishing the kinetic modeling method for dynamic FDG-PET characterization of liver inflammation, this work adds a new contribution to the methodology of dual-input kinetic modeling. We extended and validated the method of DBIF modeling using a joint optimization framework. Previously, Kudomi *et al* estimated both aortic input function and parameters of input model from multiple regional liver TACs in dynamic FDG-PET (*34*). Their method, however, is more complex for practical use with multiple steps involved. When the tracer distribution is uniform in the liver, TAC data would also be less helpful for the estimation of the many parameters in their model. Our method is simpler and can be implemented voxel-by-voxel to easily derive parametric images. The proposed method shares a similar spirit with the work by Chen and Feng (*40*) designed for modeling a different tracer ^11^C-acetate. The compartmental model used in (*40*) was irreversible while a reversible model was required in our study for FDG modeling. The major parameter of interest was *K*_*t*_ in their study, while in this study we are mainly interested in FDG K_1_. Thus, the current work has complemented existing studies to establish the optimization-derived DBIF modeling method.

Another notable aspect of this work is the validation of dual-input kinetic modeling using histopathological reference. All existing studies of dual-input kinetic modeling were only able to validate the DBIF approaches in animal studies (e.g., pigs (33,35), foxhounds (21)) or demonstrated the improvement using statistical TAC fitting quality in human patients (40). No studies had demonstrated an impact on physiological measurements by using a histopathological ground truth. In comparison, the present work provided a direct evidence on the impact of DBIF on improving association of the FDG-PET biomarker with histopathological inflammation in human patients.

There are limitations with the present study. Histopathological inflammation can be varying across different spatial locations in the liver. The statistical analysis of PET data in the study was not specific to the location of biopsy. This is because accurate biopsy location information was difficult to obtain even with ultrasound guidance and not available in this study. We chose to analyze the PET data based on the whole liver region to achieve robustness to locations and to noise in PET data. It is noted that all the studying kinetic models did not achieve a statistical significance (p>0.05) in differentiating low versus medium inflammation levels (Table 3), though the FDG K_1_ values by the optimization-derived DBIF model tended to differ in the two groups (Fig. 6C). This might be limited by the number of samples (N=14) in this study and is worth further investigation in a large number of samples.

## CONCLUSION

In this study, we examined three different kinetic models for analyzing dynamic FDG-PET data for characterizing liver inflammation. Statistical fit quality metrics and analysis of association with histopathology indicated that modeling of DBIF is crucial for accurate kinetic modeling of liver time activity curves. The optimization-derived DBIF model improved the association of FDG K_1_ with liver inflammation grades and was more appropriate than traditional single-blood input function and population-based DBIF for dynamic FDG-PET kinetic analysis in human NASH studies.

